# Cancer-associated KBTBD4 mutations induce differentiation defects and confer a unique therapeutic vulnerability

**DOI:** 10.64898/2026.03.12.711277

**Authors:** Rohit Sivaprasad, Kristijonas Žemaitis, David Linfeldt, Sudip Ghosh, Anne de Snaijer, Mattias Magnusson, Fredrik Ek, Jenny Hansson, Agatheeswaran Subramaniam

## Abstract

Epigenetic regulation governs stem cell fate and perturbations in these mechanisms often lead to tumor development. The CoREST complex, a critical regulator of neural and hematopoietic stem cell differentiation, is recurrently targeted by gain of function mutations in the ubiquitin ligase KBTBD4 in high-risk embryonal brain tumors. However, the tumorigenic potential of these mutations remains unresolved partly due to the challenges in modelling tumors that arise during early brain development. We and others recently demonstrated that small molecule UM171 mimics KBTBD4 mutations by promoting robust CoREST degradation and expansion of hematopoietic stem cells. Leveraging this mechanistic similarity, we modeled KBTBD4 mutations in hematopoietic stem and progenitor cells (HSPCs) and found that mutants induced expansion of immature stem and progenitor populations and impaired lineage differentiation. High-throughput screening identified HDAC inhibitors as specific agents that disrupt mutant KBTBD4 activity, by preventing interaction with the CoREST complex. Using our HSPC model, we demonstrated that the class I HDAC inhibitor mocetinostat alleviated differentiation defects caused by KBTBD4 mutations. Together, these findings reveal the tumorigenic mechanism of KBTBD4 mutations and uncover therapeutic vulnerabilities that may be exploited for clinical applications.

## INTRODUCTION

The ubiquitin proteasomal system tightly controls the abundance of key regulatory proteins in stem cells and the specificity of substrate recognition is achieved through a large group of E3 ubiquitin ligases. E3 ligases are often complex structures consisting of a Ring Finger Protein, a substrate receptor and an E2 enzyme to carry out the ubiquitination (1). E3 ligases recognize their respective target proteins in a highly specific manner but the specificity is altered by perturbations to the substrate recognition domains such as mutations or binding of small molecules (2, 3). Recently, we discovered that the small molecule UM171 modulates the E3 ligase CUL3-KBTBD4 to trigger a rapid polyubiquitination and degradation of chromatin associated CoREST complex (4, 5). Loss of CoREST complex promotes a strong stem cell signature and drives a robust expansion of hematopoietic stem cells (HSCs) (5).

Recurrent in-frame insertions are observed in KBTBD4 in brain tumors such as medulloblastoma and pineoblastoma (6, 7). In medulloblastoma these mutations particularly occur in group 3 and 4 tumors which are associated with poor prognosis and currently lack targeted therapies (8). More importantly, these mutations cluster to the substrate recognizing kelch domain, particularly at a hot spot spanning six amino acids (7). Remarkably, these mutants phenocopy the effect of UM171 to promote degradation of the CoREST complex and the structural basis of this similarity has been recently demonstrated (9, 10). Despite a clear molecular mechanism, the pathogenic capacity of these mutants in primary cell models and their therapeutic vulnerabilities are yet to be determined. Since medulloblastoma develops during embryogenesis, the complexity of early human brain development limits the applications of genetically engineered mouse models. In addition to this, diverse origins of subgroups become a main obstacle to model the disease (11). Therefore, developing alternate modelling approaches will be essential to gain deeper insights into the tumor biology and development of targeted therapies.

Here, we modelled KBTBD4 mutations in primary human hematopoietic stem cells (HSCs) by leveraging the mechanistic similarities of CoREST degradation in neural and blood stem cells. We demonstrate that KBTBD4 mutants promote a marked expansion of the immature HSC populations and induce severe differentiation defects. We further identified that HDAC inhibitors prevent the CoREST degradation and demonstrated that treatment with mocetinostat reversed the differentiation defects induced by KBTBD4 mutants.

## MATERIALS AND METHODS

### Umbilical cord blood isolation

Umbilical cord blood was collected from Skåne University Hospital in Lund and Malmö as well as from the Helsingborg General Hospital, all located in Sweden. The blood was spun down at 850 xg at room temperature (RT) in lymphoprep tubes (#18001, Serumwerk) for 20 mins. The mononuclear cells were collected and enriched for CD34^+^ HSPCs by magnetic bead-based isolation (#130-046-702, Miltenyi Biotech).

### Cell culture and compounds

HL60 cell cultures were grown in RPMI 1640 (#SH30605.01, Cytiva) with the addition of 10% fetal bovine serum (FBS) and 1% Pen/Strep (#15140122, Gibco). For lentiviral production, HEK293T cells were grown in DMEM (#SH30243.01, Cytiva) supplemented with 10% FBS and 1% Pen/Strep. For HSC expansion culture, StemSpan SFEM (#09650, STEMCELL™ Technologies) supplemented with stem cell factor (SCF; #300-07, PeproTech), thrombopoietin (TPO; #300-18, PeproTech), and FLT-3 Ligand (FLT3L; #300-19, PeproTech) at 100ng/mL each were used. Belinostat (#HY-10225), Vorinostat (#HY-10221), and Mocetinostat (#HY-12164) were purchased from MedChemExpress. UM171 (#72914) was purchased from STEMCELL™ Technologies. All compounds were dissolved in DMSO.

### Cloning

The KBTBD4 open reading frame was cloned into a pRRL lentiviral vector with a PGK promoter and expressing Kusabira orange as a reporter gene. The inserts were synthesized and cloned into the same vector using standard molecular cloning techniques. The procedure for cloning the ELM2-GFP wildtype (WT) and alanine substitution clones were described in our previous work (4).

### Lentiviral production and transduction

To produce lentiviral particles, HEK293T cells were transfected with a mix consisting of 10μg plasmid DNA, 7.5μg of psPAX2 and 2.5μg of pMD2.G plasmid DNA in a final volume of 2mL with Opti-MEM (#31985062, Gibco) together with 60μL of PEI (1mg/mL). The mix was vortexed and incubated at room temperature for 30 mins. Prior to transfection, the media was changed to DMEM without FBS and Pen/Strep followed by media change to complete DMEM after 4 to 6 hours. The viral supernatants were collected at 48 hours and 72 hours post-transfection and passed through a 0.45μm filter to remove the cellular debris. The viral particles were enriched either with a lenti-X concentrator (#631232, Takara) according to the manufacturer’s protocol or by ultracentrifugation at 125 000rpm for 90 mins at 4°C. For all transductions performed, the media was replaced after 16 hours to avoid toxicity from lingering viral particles.

### ELM2 degradation assay

To evaluate the degradation of various substrates by the KBTBD4 mutants, a high throughput flow cytometry based ELM2-GFP reporter assay was developed. HL60 cells were initially transduced with ELM2 domains from various candidate co-repressor proteins (RCOR1, ROCR3, and MIER3). Once the cells recovered from the transduction stress, a second round of transductions with individual KBTBD4 mutants was performed. Flow cytometry acquisition was performed at 72 and 96 hours post second transduction. ELM2 wild type and alanine substitution clones were treated with UM171 for 6 hours and the ELM2-GFP degradation efficiency was measured using flow cytometry.

### Flow cytometry

Flow cytometry analysis was performed by resuspending cells in FACS staining buffer that consists of 1X phosphate buffered saline (PBS) supplemented with 2% FBS and 2mM EDTA. 7-Aminoactinomycin D (7-AAD) was diluted 1:500 with the same buffer and was used as a dye to exclude dead cells. The acquisition was performed on BD FACSCanto, BD LSRFortessa, BD LSRFortessa X20, BD Symphony A1. For fluorescence activated cell sorting, BD FACSAria III and BD FACSDiscover S8 were used. The following antibodies were used for flow cytometry studies: CD34-PE Cy7 (#343516), CD90-FITC (#328108), EPCR-APC (#351906), CD38-PE (#356604), CD71-FITC (#555536), CD11b-APC (#101212, all from BioLegend) and CD45RA-eFluor 450 (#560362; BD Biosciences).

### Colony forming assay and liquid differentiation assay

Colony forming capacity of HSPCs were assessed in MethoCult (#H4230, STEMCELL™ Technologies) supplemented with 19% Iscove’s Modified Dulbecco’s Medium, 1% P/S and the following cytokines: SCF (#300-07, PeproTech), GM-CSF (#130-093-867, Miltenyi Biotec), IL-3 (#130-095-069, Miltenyi Biotec), EPO (#044323, Apotek) at a final concentration of 25ng/mL, 50ng/mL, 25ng/mL and 4U/mL respectively. Three hundred CD34^+^ cells were plated in a 6-well plate before incubating at standard conditions, and the colonies were counted after 13 days. Liquid differentiation assay was performed in Myelocult H5100 (#05150, STEMCELL™ Technologies) supplemented with the same cytokines as above. Flow cytometry based read outs were performed 5 days post induction of the differentiation.

### Quantitative mass spectrometry

HL60 cells were transduced with either WT KBTBD4 or P311PP mutant and collected for quantitative mass spectrometry at 72 hours post transduction. The cells were washed twice with PBS and snap frozen in dry ice. The samples were processed and analyzed at the Lund Stem Cell Center Proteomics Facility. Details about the sample processing, acquisition and analysis is added in supplementary materials.

### Small molecule screening

The screen was performed with a drug repurposing library consisting of 414 small molecules (Anticancer compound library L3000, Selleckchem). HL60 cells stably expressing RCOR1-GFP were seeded in 96-well plates at 50,000 cells per well and treated with 1µM of each compound and 200nM UM171. After 3 hours of co-treatment, cells were analyzed by flow cytometry to determine GFP levels and cell viability via 7-AAD staining, to measure RCOR1-GFP degradation rescue in live cells.

### Western blotting and Co-immunoprecipitation (Co-IP)

Cells were harvested and washed three times with ice-cold PBS prior to lysis in RIPA buffer (#10017003, Thermo Fisher Scientific) supplemented with Halt Protease Inhibitor Cocktail (#87786, Thermo Fisher Scientific) and Pierce Universal Nuclease for Cell Lysis (#12156490, Thermo Fisher Scientific). The cells together with RIPA buffer were incubated on ice for 15 mins with intermittent vortexing. The cell lysates were mixed with 2x Laemmli buffer (#161-0737, BioRad) containing Halt Protease and Phosphatase Inhibitor Cocktail (#78440, Thermo Fisher Scientific) and 5% β-mercaptoethanol. The samples were denatured at 95°C for 2 mins and run on NuPAGE electrophoresis system as per the manufacturer’s protocol. The proteins were transferred to iBlot 2 PVDF membranes (#IB24001, Thermo Fisher Scientific) and probed with the following primary antibodies: RCOR1 (#14567), KDM1A (LSD1, #2184S), HDAC2 (#5113T, all from Cell Signaling Technology), GFP (#ab290, Abcam), and Actin (#612656, BD Biosciences). All washing steps were performed 3 times for 5 mins with PBST. Blots were incubated with a HRP-conjugated secondary antibody and visualized using ECL Select HRP substrate (#12644055, Thermo Fisher Scientific). Image acquisition was performed in ChemiDoc XRS+ System according to the manufacturer’s protocol.

For Co-IP experiments, 10 million transduced HL60 cells overexpressing each alanine substitution clones were harvested and washed three times with ice-cold PBS. Cells were then lysed in ice-cold IP lysis buffer (#87788, Pierce) supplemented with Halt Protease and Phosphatase Inhibitor Cocktail, EDTA-free (100X) (#78441, Thermo Fisher Scientific), and incubated on ice for 30 minutes. Lysates were clarified by centrifugation at 18,000 × g for 15 minutes at 4°C, and the supernatant was carefully collected. In parallel, Protein A/G magnetic beads (#26162, Pierce) were blocked with 2% bovine serum albumin (BSA) in PBS for 1 hour at 4°C with rotation, followed by two washes with PBS to remove excess BSA. To reduce nonspecific binding, the collected lysates were precleared with Protein A/G beads for 30 minutes at 4°C with rotation. Beads were then separated using a magnetic stand, and the cleared lysates were incubated overnight at 4°C under constant rotation with either a control immunoglobulin G (IgG) or anti-GFP antibody (#ab290, Abcam). The following day, immune complexes were captured by incubating the antibody–lysate mixtures with fresh Protein A/G magnetic beads for 30 minutes at 4°C. Beads were subsequently washed three times with IP lysis buffer containing protease and phosphatase inhibitors, with each wash lasting 10 minutes at 4°C under rotation. To elute bound proteins, beads were resuspended in 1x Laemmli sample buffer supplemented with β-mercaptoethanol and heated at 95°C for 3 minutes. Input and immunoprecipitated fractions were then analyzed by western blotting as described above.

### Statistics

One-way ANOVA was used to compare means across multiple groups based on one categorical independent variable and two-way ANOVA was used when means were compared for multiple groups across two categorical independent variables. Results where p values ≤ 0.05 were considered significant. In the figures, p value is represented as * = ≤ 0.05, ** = ≤ 0.01, *** = ≤ 0.001, **** = ≤ 0.0001 and no asterisk for non-significant results.

## RESULTS

### KBTBD4 mutations exhibit distinct degradation profiles and induce lineage defects

To investigate the functional impact of KBTBD4 mutations, we first assessed their ability to degrade corepressor targets. We recently showed that RCOR1, RCOR3, and MIER3 containing corepressor complexes are the primary targets for degradation of UM171 induced KBTBD4 neomorphic activity (4). We chose seven commonly occurring KBTBD4 mutations and quantified the degradation profiles with a GFP based reporter system. This system consisted of individual ELM2 domains of RCOR1, RCOR3 and MIER3 tagged with GFP to monitor their levels in the cell. Co-expression of each KBTBD4 mutant with the individual ELM2-GFP constructs revealed that all insertion variants induced robust degradation of RCOR1 and RCOR3 ELM2 domains with R313PRR and P311PP being the stronger candidates (**Figure 1A, supplementary table 1**). Notably, these two mutations were observed at a higher frequency in medulloblastoma tumors (7). All the KBTBD4 mutants induced a weaker but a significant degradation of MIER3 ELM2 domain, a pattern observed previously with UM171 (4). We included one substitution mutant I310F which showed a moderate degradation of RCOR1 and no activity towards RCOR3. In summary, we successfully mapped the neomorphic activity of seven KBTBD4 mutants with the ELM2 reporter system and chose two stronger (R313PRR and P311PP) and a weaker (R312RG) candidate for the subsequent modelling studies in HSPCs.

**Figure 1:**
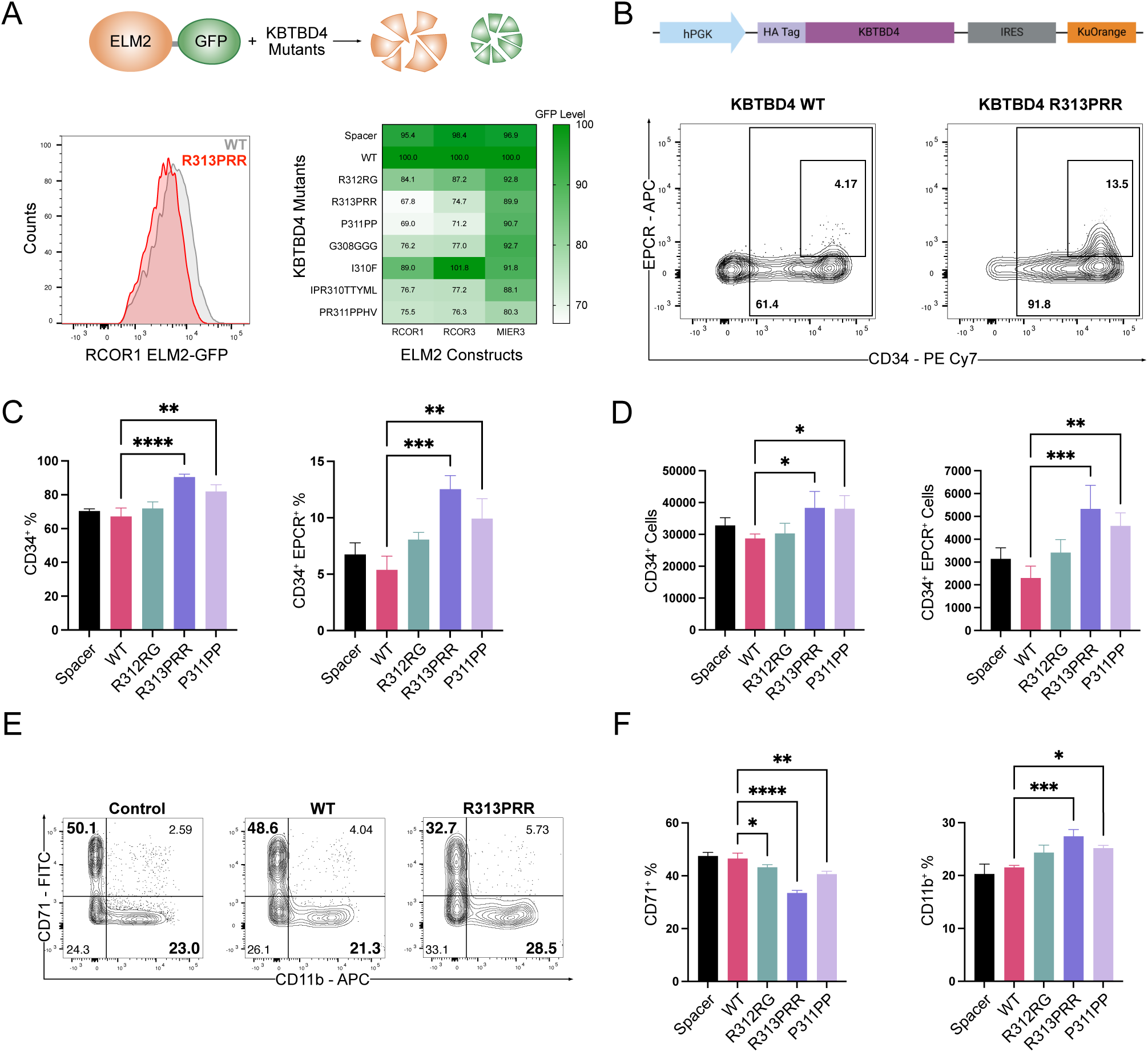
KBTBD4 mutants display distinct degradation profiles and cause differentiation defects. (A) Schematic representation of ELM2-GFP degradation induced by KBTBD4 mutants (top). FACS plot demonstrating ELM2 (RCOR1)-GFP expression in HL60 cells transduced with wild type and R313PRR mutant (left) and heat map summarizing the degradation of RCOR1, RCOR3 and MIER3 ELM2 domains by seven KBTBD4 mutants (right). Heat map was plotted with the mean of 3 replicates. (B) Stem cell enriched CD34^+^CD38^-^CD90^+^CD45RA^-^ population was FACS sorted and transduced with lentiviral vectors carrying either spacer or wildtype or mutant KBTBD4, and expanded for 7 days. Schematics of the gene construct used in this study. HA-tagged KBTBD4 constructs were expressed under the hPGK promoter, with Kusabira Orange serving as a fluorescent reporter downstream of an IRES sequence (top). FACS plots showing the expansion of progenitor (CD34^+^) and stem cells (CD34^+^EPCR^+^) cells caused by R313PRR mutant compared to the wildtype (bottom). The expansion rates of progenitor and stem cell populations for different mutants are summarized in frequencies (C) and numbers (D). Data from 3 replicates from 1 of 3 independent experiments with similar results are shown. (E) FACS plots showing the frequency of erythroid (CD71^+^) and myeloid lineages (CD11b^+^) in spacer, WT and R313PRR expressing HSPCs. (F) Summary of the lineage outputs in spacer, WT and mutants at day 5. Data from 3 replicates from 1 of 4 independent experiments with similar results are shown. Statistical analyses were performed in comparison to the WT KBTBD4.

Oncogenes are known to promote the accumulation of stem and progenitor cells in an immature state, leading to differentiation defects. Consistent with this, a previous study in medulloblastoma cell lines reported that KBTBD4 mutants activate a stem cell like gene signature (12). To assess the oncogenic potential of KBTBD4 mutants, we transduced the three selected variants in umbilical cord blood derived HSCs and measured their capacity to drive stem cell expansion. The stronger variants P311PP and R313PRR induced a significant expansion of the immature stem (CD34^+^ EPCR^+^) and progenitor (CD34^+^) cell populations (**Figure 1B-D, Supplementary figure 1A**). We observed similar results when the same variants were introduced in the umbilical cord blood derived HSPCs concluding that stronger KBTBD4 mutants indeed induce expansion of the immature cell populations (**Supplementary figure 1B-D**). Next, we assessed the differentiation potential of HSPCs transduced with KBTBD4 mutants using a liquid differentiation assay. All three mutants induced a significant reduction in erythroid lineage output, accompanied by an increase in the myeloid lineage output (**Figure 1E-F)**. To further evaluate the maturation capacity, we performed colony forming assays and found that mutant HSPCs produced fewer colonies overall with a significant reduction in erythroid colonies consistent with the results from liquid differentiation assay (**Supplementary figure 1E**). Among the variants, R313PRR induced the most severe reduction of erythroid output in both assays. Since RCOR1 loss is known to cause severe reduction in erythroid colonies, these findings suggest that the differentiation defects are primarily driven by degradation of the CoREST complex (13). In summary, modelling KBTBD4 mutants in HSPCs revealed their oncogenic properties characterized by the accumulation of immature HSPCs and impaired differentiation. Notably, the severity of these defects directly correlated with the degradation profiles, establishing degradation capacity as the key determinant of the oncogenic potential of KBTBD4 mutations.

### HDAC inhibition perturbs the interaction between KBTBD4 mutants and CoREST complex

We hypothesized that preventing CoREST degradation could alleviate the differentiation defects and serve as a promising therapeutic strategy for KBTBD4-mutant tumors. Using global proteomics, we found that the mutant P311PP induced degradation of CoREST components, including RCOR1 and LSD1, in a manner similar to UM171 (**Figure 2A, supplementary figure 2A, supplementary table 2**). To identify the drugs that perturb the neomorphic activity of KBTBD4, we took advantage of the previously established RCOR1-GFP reporter system, which exhibits a robust response to UM171 treatment (4). We screened a drug repurposing library consisting of 414 compounds and identified both broad spectrum and class I specific HDAC inhibitors as the selective hits capable of preventing RCOR1-GFP degradation (**Figure 2B)**. To validate the screening results, we selected three broad spectrum inhibitors (belinostat, vorinostat and pracinostat) and two class I specific inhibitors (romidepsin and mocetinostat) and found that all the compounds prevented RCOR1 degradation in a dose dependent manner (**Figure 2C**). Except for romidepsin, all the other compounds were well tolerated by HL60 cells highlighting their potential for further studies (**Supplementary figure 2B**). For subsequent studies, we chose belinostat as a pan HDAC inhibitor and mocetinostat as a class I specific inhibitor. Since the screening and validation studies used UM171 to mimic the neomorphic activity of mutants, we next confirmed that both belinostat and mocetinostat inhibit RCOR1 degradation induced by the P311PP KBTBD4 mutant (**Figure 2D)**. Since HDAC1/2 are integral components of the CoREST complex, HDAC inhibitors prevent the interaction between mutant KBTBD4 and CoREST, consistent with recent structural studies that solved the structural basis of these interactions (9, 10). Previously, we identified critical amino acids in the N-terminus region of ELM2 domain that dictate RCOR1 degradation (4). Given that HDAC2 is the primary target of KBTBD4 neomorphic activity, we asked whether these residues are important for interaction with HDAC2. Remarkably, either deletion or alanine substitution of individual critical amino acids within the RCOR1 ELM2 domain abrogated the interaction with HDAC2 revealing the role of this interaction in promoting CoREST degradation (**Figure 2E, supplementary table 3)**. In summary, our results demonstrated that HDAC inhibitors disrupt the interaction between KBTBD4 mutants and CoREST complex, revealing a distinct therapeutic vulnerability that could be exploited to target KBTBD4-driven tumors (**Figure 2F)**.

**Figure 2:**
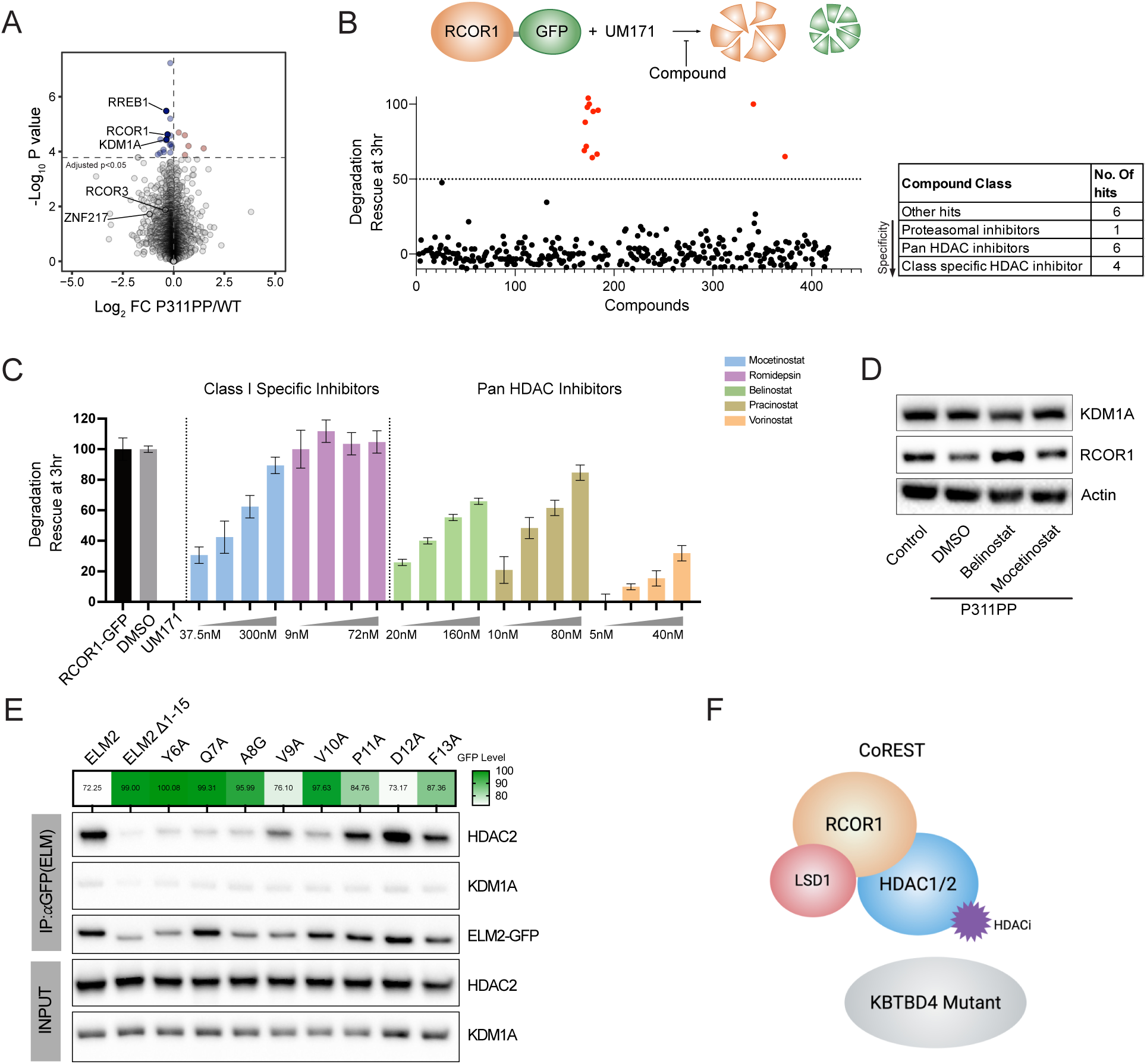
HDAC inhibitors perturb the interaction between KBTBD4 mutants and CoREST complex. (A) Volcano plot showing global protein expression in HL60 cells transduced with P311PP mutant. Blue and red dots show significantly decreased or increased proteins (adjusted p-value<0.05). CoREST components RREB1, RCOR1, KDM1A, RCOR3, and ZNF217 are highlighted in the volcano plot. The global proteome data was plotted after excluding PCDHGA11. (B) High throughput compound screening in HL60 cells stably expressing RCOR1-GFP treated with UM171 (200nM). Schematic of the screening rationale (top panel). The compounds were added at a final concentration of 1µM, and flow analysis was performed after 3 hours post treatment to measure the relative GFP expression (bottom panel). Red dots show the hits from the screen and the table on right-side shows the functional classification of the hits. (C) A dose-titration experiment showing the rescue of RCOR1-GFP by two class I specific HDAC inhibitors (mocetinostat and romidepsin) and three Pan HDAC inhibitors (belinostat, pracinostat and vorinostat). GFP mean fluorescence intensity of DMSO treatment was used as controls. Data from 3 replicates from 1 of 2 independent experiments with similar results are shown. (D) RCOR1 protein levels in P311PP expressing HL60 cells either treated with DMSO, belinostat (320nM) or mocetinostat (300nM) for 3 hours. (E) RCOR1 ELM2-GFP clones were expressed in HL60 cells and analyzed for the interaction with HDAC2 through immunoprecipitation. The degradation profiles of the corresponding alanine substitution clones to UM171 treatment are represented as a heat map. (F) Schematic representation of the mechanistic basis of HDAC inhibitors in preventing the CoREST degradation.

### Mocetinostat alleviates lineage differentiation defects induced by KBTBD4 mutants

Given that HDACs are integral components of multiple chromatin remodeling complexes, we investigated whether pharmacological inhibition of HDACs could mitigate the effects of KBTBD4 mutants. To address this, we treated HSPCs expressing KBTBD4 mutants with belinostat and mocetinostat and quantified the lineage output. Strikingly, mocetinostat treatment has effectively alleviated the lineage defects caused by all three mutants with no detectable impact on spacer and wildtype controls highlighting the specificity towards KBTBD4 mutant cells (**Figure 3A-B**). While belinostat rescued the erythroid defects, it also reduced myeloid output in both mutant and control HSPCs indicating that pan-HDAC inhibition introduces off-target effects. Importantly, both compounds were tolerated by HSPCs with no significant effect on viability but with a significant reduction in the proliferation rates (**Supplementary figure 3A-B**). We further tested these two compounds in unmanipulated HSPCs and found that belinostat treatment indeed induced myeloid lineage defects while mocetinostat was tolerated (**Supplementary figure 3C**). To test the broader applicability of mocetinostat across the most common KBTBD4 mutants we used our ELM2-GFP model and demonstrated that mocetinostat indeed perturbs the neomorphic activity of all seven mutants (**Figure 3C)**. In summary, we conclude class I HDAC inhibition with mocetinostat as a potential targeted therapy for KBTBD4-driven tumors.

**Figure 3:**
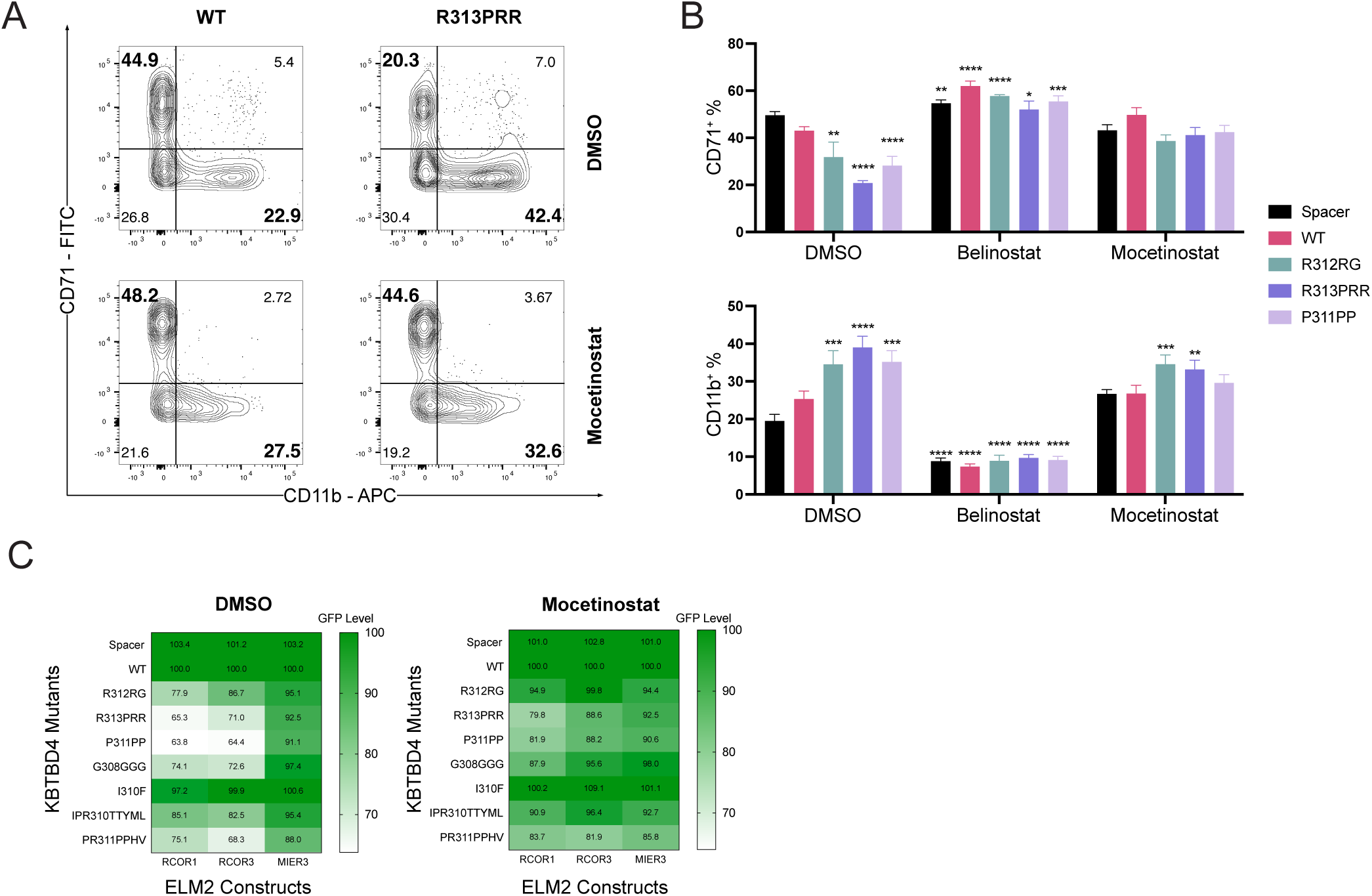
Mocetinostat alleviates lineage defects caused by KBTBD4 mutants. (A) FACS plots showing lineage output in WT and R313PRR expressing HSPCs treated with either DMSO or mocetinostat (250nM) for 5 days. (B) Summary of the lineage output for DMSO, belinostat (125nM) and mocetinostat (250nM) treatments. Data from 3 replicates from 1 of 4 independent experiments with similar results are shown. (C) Heat map demonstrating ELM2-GFP expression profiles in HL60 cells overexpressing KBTBD4 mutants treated with DMSO or mocetinostat (75nM) for 24 hours. Data analyzed at 96 hours post transduction. Statistical analyses were performed in comparison to the WT KBTBD4 treated with DMSO.

## DISCUSSION

We demonstrate the oncogenic properties of cancer-associated KBTBD4 mutants using human umbilical cord blood-derived HSPCs. Based on our findings, we propose class I HDAC inhibition as a viable therapeutic strategy to counteract the effects of KBTBD4 mutants.

Though the mutational landscapes of group 3 and 4 medulloblastoma tumors are well characterized, divergence in the early development between human and mouse makes it harder to model these mutations in vivo (11). The lack of suitable in vitro models for these high-risk tumors further limits the identification of new therapeutic opportunities. The mechanistic mimicry between HSC supporting molecule UM171 and KBTBD4 mutations converges on the CoREST complex and highlights a similar role of CoREST in neural and hematopoietic stem cell differentiation. This makes HSCs an attractive system to model the oncogenic properties of KBTBD4 mutations. Supporting this connection, activation of hematopoietic transcription factors GFI1 and GFI1B is observed as main driver events in group 3 and 4 medulloblastoma suggesting deeper biological similarities between these two systems (14).

The CoREST complex plays an essential role in the differentiation of hematopoietic and neural stem cells and the current evidence indicates that it acts as a tumor suppressor as the loss leads to medulloblastoma development. While KBTBD4 mutations degrade CoREST, GFI1/1B likely displaces it from the native targets, both converging on CoREST inactivation (15). Further studies are required to delineate the downstream targets of CoREST and to establish their functional relevance in medulloblastoma development.

HDAC inhibitors have been tested as therapeutic agents for targeting CNS tumors, nevertheless the lack of mechanism-based subgroup stratification has emerged as a major limitation to assess their therapeutic benefit (16). Our study supports the use of class I HDAC inhibitors for KBTBD4-driven medulloblastoma, however further in vivo studies with patient-derived xenografts or in the relevant organoid models will strengthen these findings. In conclusion, we modelled the oncogenic properties of KBTBD4 mutants and identified a unique therapeutic vulnerability for targeting medulloblastoma tumors.

## ACKNOWLEDGEMENTS

We would like to thank the staff at FACS core, Vector core and Proteomics facilities at Lund Stem Cell Center for their excellent support. The compound library was provided by the Chemical Biology Consortium Sweden (CBCS) Lund node, a research infrastructure funded by the Swedish Research Council (dr.nr. 2021-00179)), and MultiPark, a strategic research area at Lund and Gothenburg University. This work was funded by grants from the Swedish Cancer Foundation, Swedish Childhood Cancer Foundation, Gunnar Nilsson’s Cancerstiftelse, Åke Wibergs Stiftelse, Fru Berta Kamprad’s Stiftelse and Royal physiographic society of Lund to A.S. K.Z. is supported by a postdoctoral grant from Swedish Childhood Cancer Foundation.

## AUTHOR CONTRIBUTIONS

R.S. and A.S. planned the study, designed the experiments and analyzed the results. R.S., K.Ž., D.L., A.D.S. performed the studies and analyzed the results. S.G. performed mass spectrometry and analyzed the data together with J.H. F.E. and M.M. helped with the compound library preparations for the screening assay. R.S. and A.S. wrote the manuscript. A.S. conceived the study.

## DECLARATION OF INTERESTS

The authors declare no competing interests.

## DATA AVAILABILITY

The datasets generated in this study can be provided upon request by the corresponding author.

## Notes

### Competing Interest Statement

The authors have declared no competing interest.

