## Supplementary file for "Cancer-associated KBTBD4 mutations induce differentiation defects and confer a unique therapeutic vulnerability"

### Supplementary methods

#### Sample preparation for cellular proteome analysis

HL60 cell lines transduced with over-expression constructs of either wildtype or P311PP mutant were collected in ice-cold phosphate buffer saline solution (PBS), centrifuged, and stored as dry pellets at -80 °C until further use. Pellets corresponding to 500,000 cells were solubilized in 50 mM ammonium bicarbonate (Sigma) with 0.1% RapiGest (Waters) to denature proteins and shaken on a thermomixer (Eppendorf) at 400 rpm for 15 minutes decreasing the temperature from 80°C to 56°C, followed by reduction of disulfide bonds with 0.1M dithiothreitol at 56°C, cysteine alkylation with 0.2 M iodoacetamide at room temperature in the dark, and digestion overnight at 37°C with sequencing grade modified trypsin (enzyme: protein ratio 1:50, Promega). Digested peptides were acidified with 10% trifluoroacetic acid (TFA) and RapiGest was precipitated by incubation at 37°C followed by centrifugation. Peptides were desalted with SepPak C18 columns (Waters), dried by vacuum centrifugation, and resuspended in MS loading buffer prior to LC-MS analysis.

#### Liquid chromatography and mass spectrometry (LC-MS)

LC-MS analyses were carried out on an Orbitrap Exploris 480 MS instrument with a reverse phase UltiMate 3000 UHPLC system via an EASY-Spray ion source equipped with FAIMS Pro (all Thermo Fisher Scientific). Peptide samples were analyzed by data-independent acquisition (DIA). Digested peptides were loaded onto a trap cartridge (Acclaim PepMap C18, 5 mm particle size, 0.3 mm inner diameter x 5 mm length, Thermo Fisher Scientific) and separated on an EASY-Spray analytical column (2 mm particle size, 75 mm inner diameter x 500 mm length, Thermo Fisher Scientific). Each sample was injected once and eluted with a linear gradient ranging from 2-19% Solvent B (0.1%FA in 80% ACN) over 80 min, 19-41% B over 40 min, 41-90% B over 5 min and held at 90-95% B for 5 min at a constant flow rate of 300 nl/min at 45°C. FAIMS compensation voltages (CV) were set to -45 and -60. The spray voltage was set at 2.1 kV and the ion transfer tube temperature was set at 275°C. For DIA analysis, peptides were analyzed with one full scan (350–1,400 m/z, R = 120,000) at a normalized AGC target of 300%, followed by 38 DIA MS/MS scans (350–1,050 m/z) in HCD mode (isolation window 22 m/z, 1 m/z window overlap, normalized collision energy 27%), with fragments detected in the Orbitrap (R = 15,000). All data were acquired in positive polarity and MS/MS were acquired in centroid mode.

**MS raw data processing and Statistical analysis**

The MS data were searched with 'directDIA' in Spectronaut (version 18, Biognosys AG) against the human SwissProt reference proteome and commonly used contaminants. Searches used carbamidomethylation as fixed modification, and methionine oxidation as well as protein N-terminal acetylation as variable modifications. The Trypsin/P proteolytic cleavage rule was used, permitting a maximum of 2 missed cleavages and a minimum peptide length of 7 amino acids. 'Cross run normalization' was enabled with Normalization Strategy set to 'local normalization' based on rows with 'Identified in All Runs (Complete)'. Data filtering was set to Q-value and the Q-value thresholds were set to 0.01 at PSM, peptide, and protein levels. Protein quantification and statistical analysis were performed with Msstats (1) (version 4.17.1). Contaminants were filtered and features were converted to MSStats format for downstream processing. Uninformative features were removed and missing values were imputed with the 'MBimpute' function within MSStats. Statistical analysis was performed with MSStats 'group comparison'. Differentially expressed proteins were considered with adjusted p-value of less than 0.05 between groups for each comparison.


#### Supplementary figure legends

**Supplementary Figure 1.** (A) Transduction efficiency of KBTBD4 mutants in HSC enriched $CD34^{+}CD38^{-}CD90^{+}CD45RA^{-}$ population and (B) the $CD34^{+}$ bulk stem and progenitor cells. The bulk $CD34^{+}$ cells were transduced with KBTBD4 mutants, and the frequency (C) and numbers (D) of progenitor ($CD34^{+}$) and stem cells ($CD34^{+}$ $EPCR^{+}$) were measured at 7 days post transduction. (E) Colony forming capacity of stem and progenitor cells transduced KBTBD4 mutants. The HSPCs were transduced and expanded for 7 days and plated for 13 days to induce colony formation. Data are shown as the average of 3 replicates from 1 of 3 independent experiments with similar results. Statistical analyses were performed in comparison to the WT KBTBD4.

**Supplementary Figure 2.** (A) A comparison of degradome profiles between the P311PP mutant and molecular glue UM171 as quantified in mass spectrometry ('*' denotes adjusted p-value < 0.05) (B) Viability of HL60 RCOR1-GFP cells with the treatment of two class I specific HDAC inhibitors (mocetinostat and romidepsin) and three Pan HDAC inhibitors (belinostat, pracinostat and vorinostat). Cells were treated with the inhibitors for 3 days and the viability range was normalized to DMSO controls. Data from 3 replicates from 1 of 2 independent experiments with similar results are shown.

**Supplementary Figure 3.** (A) Viability of the modelled HSPCs treated with belinostat (125nM) and mocetinostat (250nM) for 5 days. (B) Erythroid ($CD71^{+}$) and myeloid ($CD11b^{+}$) lineage numbers in HSPCs expressing either spacer, wild type or KBTBD4 mutants and treated with either DMSO, belinostat (125nM) or mocetinostat (250nM) for 5 days. (C) lineage output in HSPCs treated with belinostat (125nM) and mocetinostat (250nM) for 5 days. CD71-Erythroid marker; CD11b-Myeloid marker.

### Supplementary figures

#### Supplementary Fig 1

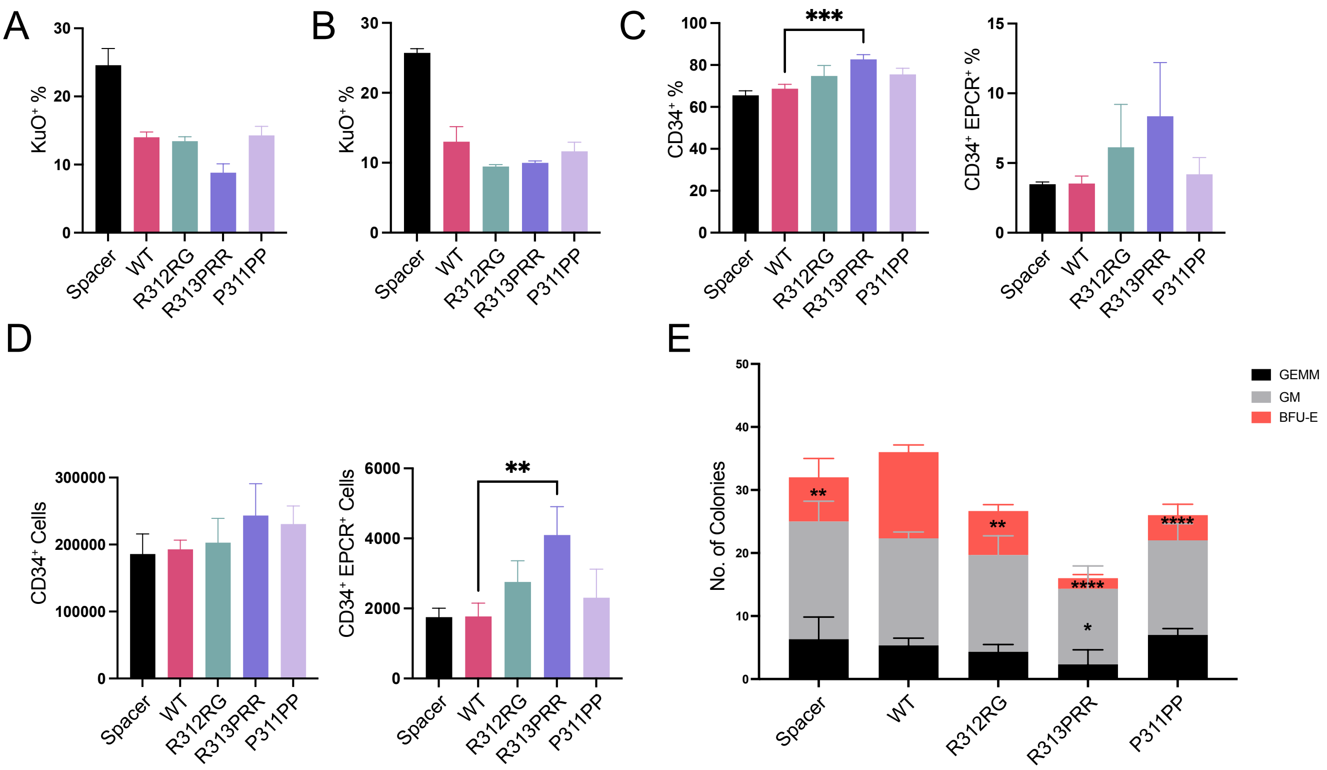


#### Supplementary Fig 2

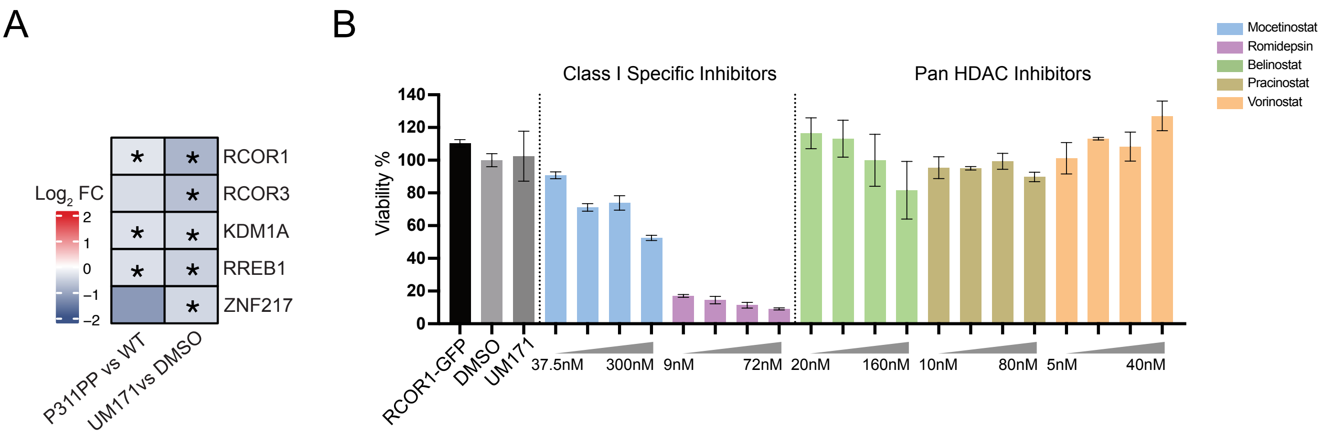

### Supplementary Fig 3

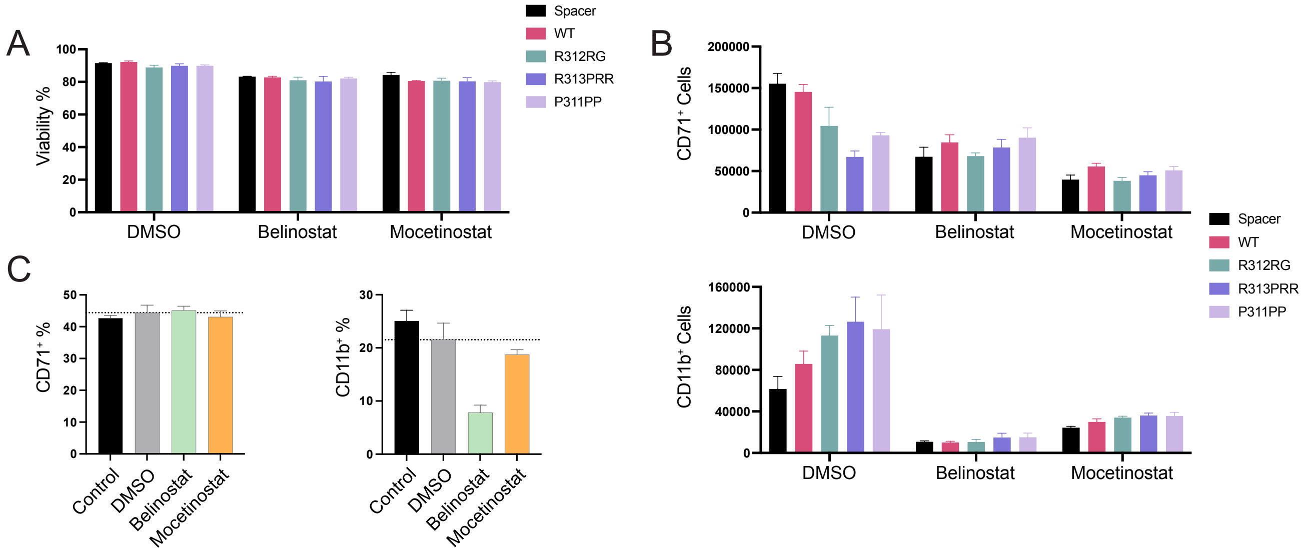

#### Supplementary table legends

**Supplementary Table 1.** ELM2-GFP mean fluorescent intensity (MFI) values for RCOR1, RCOR3 and MIER3 ELM2 domains when co-expressed with individual KBTBD4 mutants.

**Supplementary Table 2.** Quantitative proteomic analysis in HL60 cells transduced with P311PP mutant in comparison to the wild type KBTBD4.

**Supplementary Table 3.** Mutant ELM2 clones used in alanine scan. Amino acids in RCOR1 ELM2 domain were substituted with either alanine or glycine (where WT alanine was present).

#### Supplementary tables

**Supplementary Table 3**

| Position within RCOR1 ELM2 Domain | Wildtype Amino acid | Substituted Amino acid |
|---|---|---|
| 6 | Y | A |
| 7 | Q | A |
| 8 | A | G |
| 9 | V | A |
| 10 | V | A |
| 11 | P | A |
| 12 | D | A |
| 13 | F | A |
